# Detecting cell-type-specific allelic expression imbalance by integrative analysis of bulk and single-cell RNA sequencing data

**DOI:** 10.1101/2020.08.26.267815

**Authors:** Jiaxin Fan, Xuran Wang, Rui Xiao, Mingyao Li

## Abstract

Allelic expression imbalance (AEI), quantified by the relative expression of two alleles of a gene in a diploid organism, can help explain phenotypic variations among individuals. Traditional methods detect AEI using bulk RNA sequencing (RNA-seq) data, a data type that averages out cell-to-cell heterogeneity in gene expression across cell types. Since the patterns of AEI may vary across different cell types, it is desirable to study AEI in a cell-type-specific manner. Although this can be achieved by single-cell RNA sequencing (scRNA-seq), it requires full-length transcript to be sequenced in single cells of a large number of individuals, which are still cost prohibitive to generate. To overcome this limitation and utilize the vast amount of existing disease relevant bulk tissue RNA-seq data, we developed BSCET, which enables the characterization of cell-type-specific AEI in bulk RNA-seq data by integrating cell type composition information inferred from a small set of scRNA-seq samples, possibly obtained from an external dataset. By modeling covariate effect, BSCET can also detect genes whose cell-type-specific AEI are associated with clinical factors. Through extensive benchmark evaluations, we show that BSCET correctly detected genes with cell-type-specific AEI and differential AEI between healthy and diseased samples using bulk RNA-seq data. BSCET also uncovered cell-type-specific AEIs that were missed in bulk data analysis when the directions of AEI are opposite in different cell types. We further applied BSCET to two pancreatic islet bulk RNA-seq datasets, and detected genes showing cell-type-specific AEI that are related to the progression of type 2 diabetes. Since bulk RNA-seq data are easily accessible, BSCET provided a convenient tool to integrate information from scRNA-seq data to gain insight on AEI with cell type resolution. Results from such analysis will advance our understanding of cell type contributions in human diseases.

**Author Summary:** Detection of allelic expression imbalance (AEI), a phenomenon where the two alleles of a gene differ in their expression magnitude, is a key step towards the understanding of phenotypic variations among individuals. Existing methods detect AEI use bulk RNA sequencing (RNA-seq) data and ignore AEI variations among different cell types. Although single-cell RNA sequencing (scRNA-seq) has enabled the characterization of cell-to-cell heterogeneity in gene expression, the high costs have limited its application in AEI analysis. To overcome this limitation, we developed BSCET to characterize cell-type-specific AEI using the widely available bulk RNA-seq data by integrating cell-type composition information inferred from scRNA-seq samples. Since the degree of AEI may vary with disease phenotypes, we further extended BSCET to detect genes whose cell-type-specific AEIs are associated with clinical factors. Through extensive benchmark evaluations and analyses of two pancreatic islet bulk RNA-seq datasets, we demonstrated BSCET’s ability to refine bulk-level AEI to cell-type resolution, and to identify genes whose cell-type-specific AEIs are associated with the progression of type 2 diabetes. With the vast amount of easily accessible bulk RNA-seq data, we believe BSCET will be a valuable tool for elucidating cell type contributions in human diseases.

## Introduction

Allelic expression imbalance (AEI) refers to the phenomenon where the expression between the paternal and maternal alleles of a gene differs in their magnitude in a diploid individual^1^. In the presence of *cis*-regulatory effect, the expression increasing allele at a *cis*-regulatory polymorphism can lead to higher expression of one allele compared to the other. Such allelic imbalance in gene expression may associate with phenotypic variations among individuals and contribute to human disease. Since AEI measures the expression of two alleles expressed in the same cellular environment, evidence of AEI can rule out the influence of *tran*-regulatory variants, and thus is ideal for detecting *cis*-regulatory effects and pinpointing causal variants for disease^2^.

Traditionally, AEI is studied by bulk RNA sequencing (RNA-seq) in which the allelic expression differences at heterozygous exonic single nucleotide polymorphisms (SNPs) are characterized by allele-specific read counts^3–5^. However, solid tissue is typically composed of cells that originate from diverse cell types, and bulk RNA-seq can only measure the average expression across all cells in a bulk tissue sample. Previous studies have shown that gene expression is often altered in a cell-type-specific manner, and it is possible that only certain cell types are responsible for phenotypic changes^6, 7^. To gain further insight into the *cis*-regulatory effect, it is necessary to characterize AEI with cell type resolution.

Recent advances in single-cell RNA sequencing (scRNA-seq) technologies have enabled researchers to characterize individual cells and study gene expression with cell type resolution even for rare cell populations^8^. However, to study the cell-type-specific effect of AEI across individuals, full-length transcript sequencing in single cells across a large number of individuals is needed. Given the current cost of scRNA-seq, it is still cost prohibitive to generate such data. Since a large amount of bulk RNA-seq data in disease relevant tissues have already been generated in previous studies, it is desirable to characterize cell-type-specific AEI using these existing data by leveraging cell-type information provided by scRNA-seq.

The goal of this paper is to develop a two-step regression-based procedure that allows us to integrate **<u>B</u>**ulk and **<u>S</u>**ingle-cell RNA-seq data to detect **<u>CE</u>**ll-**<u>T</u>**ype-specific allelic expression imbalance (BSCET). First, we perform cell-type deconvolution analysis to infer cell type composition in bulk RNA-seq data. Second, given estimated cell type proportions in bulk RNA-seq samples, we test cell-type-specific AEI using allele-specific bulk RNA-seq read counts. Since the degree of AEI may vary with disease phenotypes, we further extend BSCET to incorporate clinical factors such as disease status to infer covariate effects on cell-type-specific AEI. Through comprehensive benchmark evaluations and real data applications to two pancreatic islet bulk RNA-seq datasets, we show that BSCET is able to detect cell-type-specific AEI and differential cell-type-specific AEI between healthy and diseased samples.

## Results

### Overview of methods and evaluation

The primary goal of BSCET is to detect cell-type-specific AEI across individuals using bulk RNA-seq data. It takes two datasets as input, a bulk RNA-seq dataset, which is used to detect cell-type-specific AEI, and a scRNA-seq dataset that is used as a reference to infer cell type compositions in the bulk data. After cell type compositions for all bulk RNA-seq samples are inferred, for each transcribed SNP, BSCET then aggregates allele-specific read counts from the bulk RNA-seq data across individuals to model cell-type-specific expression difference between two alleles by linear regression, and tests for the evidence of cell-type-specific AEI. For a SNP with cell-type-specific AEI, it might be of interest to further investigate if its cell-type-specific AEI is affected by any clinical factors. This can be achieved by including covariates in the regression. Specifically, BSCET models the difference in expression between the two transcribed alleles of the SNP over cell type compositions and includes interactions between cell type compositions and the covariate of interest. By testing the interaction term, BSCET assesses whether this covariate alters the cell-type-specific AEI in the corresponding cell type. An overview of BSCET is shown in **Fig 1**, and details can be found in **Methods**.

**Fig 1.**
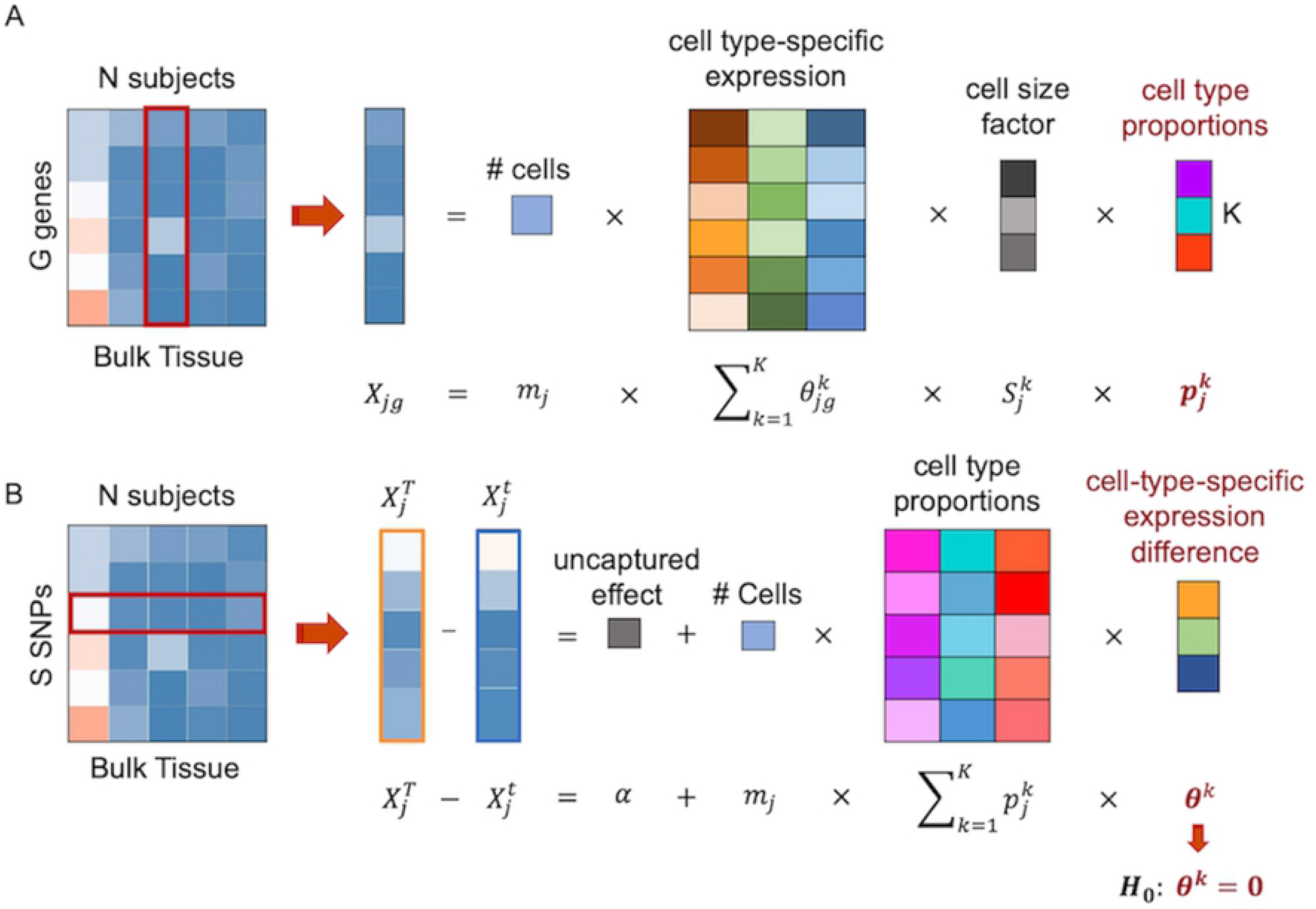
Overview of BSCET. The method contains two steps. **(A)** First, we employ the deconvolution method, MuSiC^11^, to infer the cell type proportion, 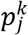, of the bulk tissue for each individual *j* of cell type *k* by incorporating the cell-type-specific expression information, 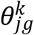, and cell size factor, 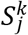, obtained from scRNA-seq data. **(B)** Second, at each transcribed SNP (*tSNP*) site, we align the *tSNP* alleles with respect to its reference and alternative alleles, T and t, across individuals, and model the expression difference between two allele-specific read counts, 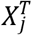 and 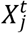, over the estimated cell type compositions, 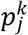, through a linear model with an intercept term *α* to capture the information not explained by the selected *K* cell types, where both *m_j_* and 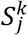 are assumed known. The difference of population-level relative abundance between two transcribed alleles, *θ^k^*, can therefore be inferred for each cell type. By testing for *H*_0_: *θ^k^* = 0, we can detect cell-type-specific allelic imbalance across samples.

We performed benchmark simulations to evaluate the performance of BSCET. The benchmark bulk RNA-seq data were generated based on a human pancreatic islet scRNA-seq dataset^9^. For the simulated benchmark data, the cell type proportions and AEI levels for each cell type are known, which allow us to evaluate the performance of BSCET. The benchmark simulation procedure was shown in **Fig 2** and details were described in **Methods**. As a comparison, we also analyzed the benchmark data using a traditional bulk AEI detection method that ignores cell-type-specific effect. In this analysis, the allelic read counts were modeled using a generalized linear model (GLM) for binomial data with logit link function. The relative expression of the reference allele over total read counts was modeled across individuals using an intercept only model, and evidence of AEI was assessed by testing whether the intercept is significantly different from zero.

**Fig 2.**
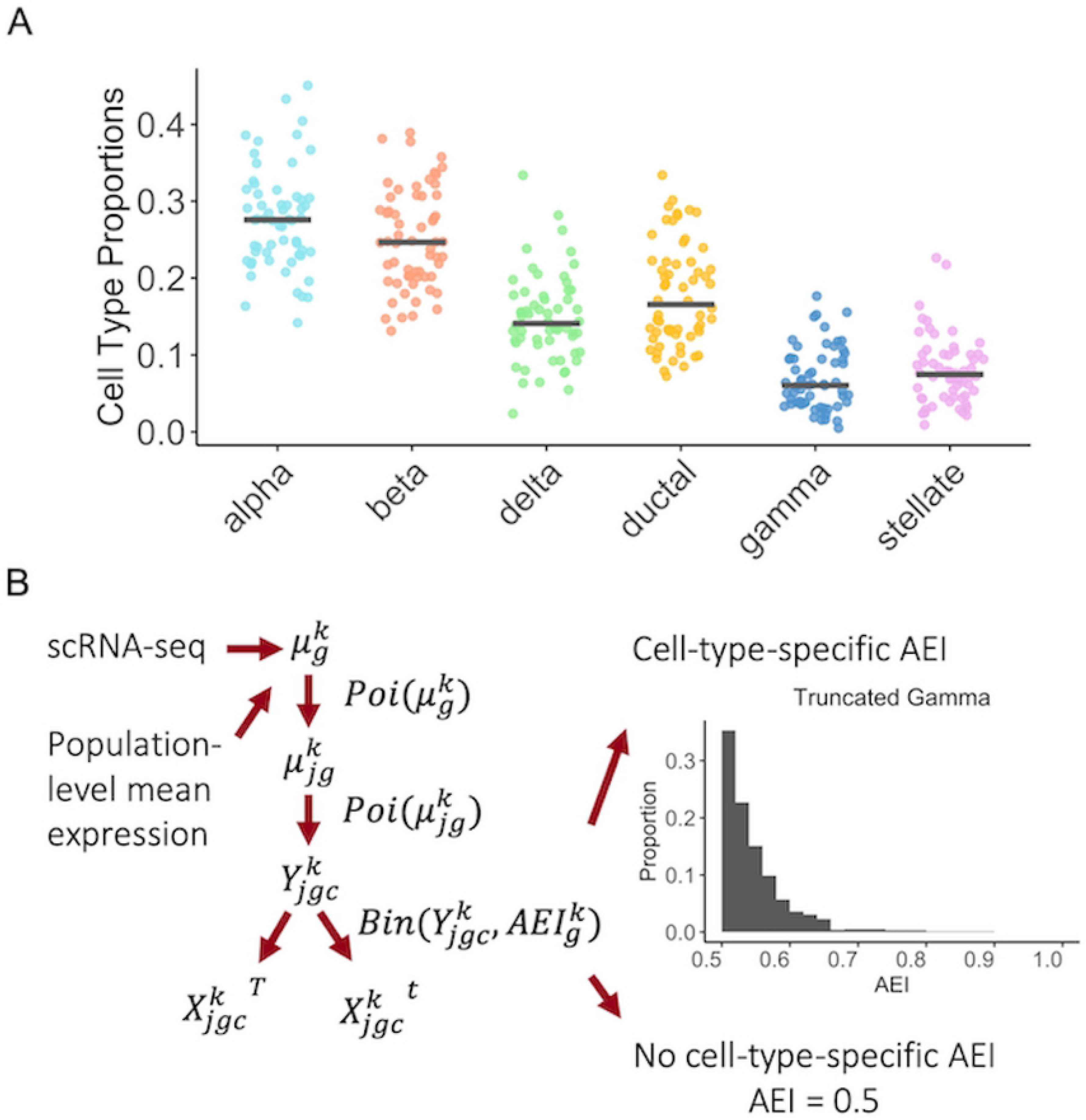
Benchmark evaluation data generation scheme. **(A)** Cell type compositions assumed for the artificial bulk RNA-deq data. For each cell type, the solid line indicates the average cell type proportions across samples, which was inferred from the single cell data^9^. **(B)** Data generation scheme for obtaining the artificial allele-specific bulk RNA-seq data. Given gene *g*, for each cell type *k*, we first inferred the mean expression level from the single cell data^9^. Based on the sample mean, we next sampled the subject-specific mean expression through Poisson distribution, and used another layer of Poisson sampling to obtain the total read count of each cell *c*. The total expression was then split into two allele-specific read counts through a binomial distribution with the probability equals to the cell-type-specific AEI. For each gene *g*, we generated data only for a single SNP, and the level of AEI equaled to 0.5 for cell types with no allelic imbalance and was generated from a truncated gamma distribution for cell types with AEI. The artificial bulk RNA-seq data was obtained by summing up allelic read counts across all cells.

### Detection of cell-type-specific AEI when one cell type has AEI

The simulated benchmark bulk RNA-seq data include 60 individuals and the data were generated with known cell type proportions and cell-type-specific AEI levels. We considered two scenarios: first, we assumed only the major cell type, had AEI; and second, we assumed both the major and a non-major cell type, had AEI. For each gene, the major cell type was selected as the cell type with the highest mean expression among all cell types, and a non-major cell type was selected as the one with median mean expression in the original scRNA-seq data in which the benchmark bulk RNA-seq data were sampled from. Under each scenario, the model performance was evaluated using both the true and “estimated” cell type proportions, where the “estimated” proportions were obtained by adding random noise to the true proportions to reflect estimation uncertainty. More specifically, for individual *j* of cell type *k*, the “estimated” proportion, 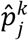, was set as 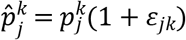, where 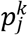 is the true proportion and the random noise, *ε*_*jk*_, was sampled from *N*(0,0.2^2^).

We analyzed 11,300 SNPs in total and evaluated the type I error on cell types that did not have cell-type-specific AEI. Under the first scenario where only the major cell type had AEI, when using true cell type proportions as input for BSCET, the false positives were under control at the 0.05 significance level (**Fig 3A**). We correctly identified 2,651 SNPs showing cell-type-specific AEI after false discovery rate (FDR) adjustment for multiple testing. The overall detection power was higher for alpha, beta, and ductal cells than delta and gamma cells, which is not surprising since alpha, beta and ductal cells are the common cell types in islets^10, 11^. Interestingly, despite the low proportion in bulk tissue, stellate cells had the highest detection power among all cell types, likely due to its highest expression level among the six cell types in the scRNA-seq data^9^. As a comparison, the traditional GLM method for bulk data analysis detected 2,813 SNPs with AEI, among which 1,662 overlapped with the SNPs detected by BSCET. We found that SNPs detected only by the bulk GLM method had a higher expression level on average (mean reads of 481.7 with interquartile range (IQR) 166.9-583.3) compared to those detected only by BSCET (mean reads of 144.2 with IQR 55.4-183.3) (**S1 Fig**). As the bulk detection method is sensitive to high read counts, whereas BSCET deconvolves the read counts into each cell type, it is not surprising that the bulk method is more powerful than BSCET when sequencing depth is high. However, the goal of BSCET is not to detect more genes than the bulk GLM method, but rather to elucidate what cell type is driving the evidence of AEI. Nevertheless, for SNPs that are not highly expressed at the bulk level, BSCET correctly detected some of them with cell-type-specific AEI while the bulk GLM method failed to detect. For most of these SNPs, the major cell type does not have high cell type proportions, thus the bulk GLM method has limited detection power as the AEI signals were diluted due to other more common cell types that do not have AEI.

**Fig 3.**
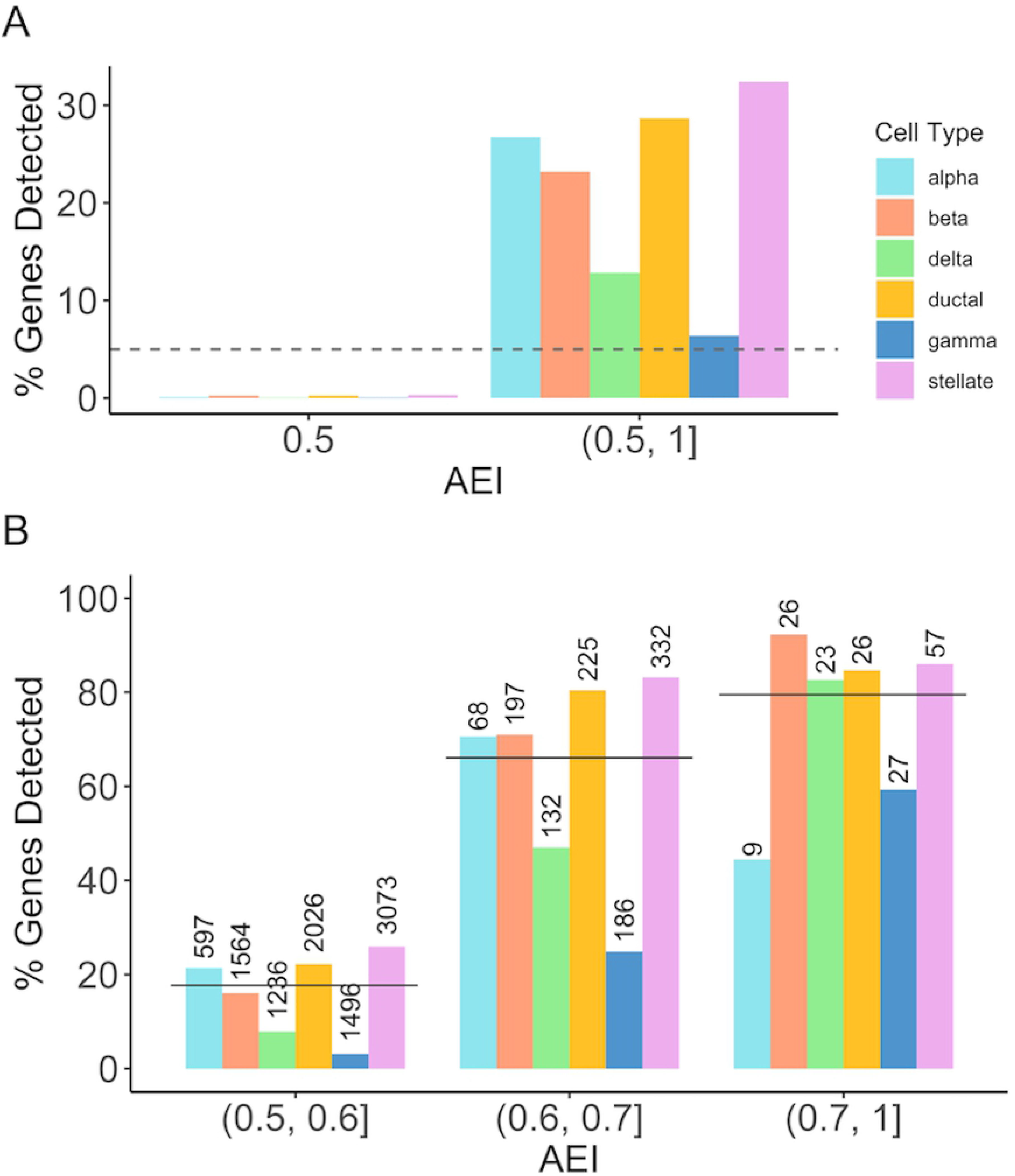
Benchmark evaluation for cell-type-specific AEI detection assuming one cell type with AEI. We evaluated the performance of BSCET assuming the major cell type for each SNP, i.e., cell type with the highest mean expression in scRNA-seq data^9^, had cell-type-specific AEI using true cell type proportions. **(A)** Type I error rate and power evaluated at the cell type resolution under significance level *α* = 0.05 (dashed line). **(B)** Detection power separated by the cell type and true level of AEI. The number above each bar indicates the total number of genes with cell-type-specific AEI within the category. The solid line indicates the overall power, i.e., across all cell types, at each level of AEI.

We further evaluated the performance of BSCET at different AEI levels (**Fig 3B**). As expected, BSCET detected more genes as the true AEI level increased, and the power reached 80% when the AEI was as large as 0.7. Similarly, the power increased with AEI level for most cell types, except for alpha cells, which was likely due to the relatively small number of SNPs in the [0.7, 1) AEI category. Further, at different AEI levels, alpha, beta, ductal and stellate cells had higher detection power in general than delta and gamma cells, and the difference became smaller as the underlying true AEI increased. BSCET performed the best in stellate cells when the AEI effect was low to moderate, indicating that it is able to detect small AEI effect even for rare cell types, as long as the expression level is relatively high.

Next, we evaluated the performance of BSCET using “estimated” cell type proportions. Again, the type I error was well controlled. Among the 11,300 tested SNPs, 2,276 were detected to have cell-type-specific AEI after FDR adjustment. The correlation of p-values obtained using “estimated” and true cell type proportions across all tested SNPs was shown in **S2A Fig**. As expected, the power was slightly lower than that when the true cell type compositions were used. However, the SNP-level p-values obtained using the “estimated” proportions were highly correlated with those obtained using the true proportions (Pearson’s correlation coefficient R = 0.94, **S2B Fig**). Further, we observed a similar increasing pattern in power over the true AEI levels as well as similar power differences between cell types, indicating that BSCET is robust to estimation uncertainty in cell type proportions (**S2C Fig**).

### Detection of cell-type-specific AEI when two cell types have AEI

We also considered the scenario in which both the major cell type and a non-major cell type had AEI. Rather than letting all SNPs had AEI for both cell types in the same direction, we assumed the effect was in opposite directions for 30% of the 11,300 tested SNPs. For these 30% of the SNPs, the AEI effects in the two cell types would cancel each other out when averaging across all cells, thus the bulk detection method might fail to detect evidence of AEI. Assuming the true cell type compositions were known, among the 7,873 (70%) tested SNPs with AEI in the same direction, BSCET correctly uncovered 1,920 for the major cell type and 1,268 for the non-major cell type, with type I error well controlled at the 0.05 significance level. For the major cell type, the results were similar to those obtained under the first scenario, indicating that the performance of BSCET was not influenced much when more than one cell type had AEI. For the non-major cell type, due to its lower expression level than the major cell type, the detection power, as expected, was lower than that for the major cell type. Nevertheless, we still had around 80% power when the underlying AEI was large, e.g., over 0.7, and the power difference between the major and non-major cell types decreased as the AEI level increased. Among the 3,427 (30%) tested SNPs with opposite AEI directions, we correctly recovered the direction of cell-type-specific AEI for 2,330 SNPs (68%), and identified 837 with AEI for the major cell type and 502 for the non-major cell type after FDR adjustment. For both the major and non-major cell types, the patterns of type I error and power are consistent with those observed in **Fig 4A**, suggesting that BSCET was reliable for detecting cell-type-specific effect even when the two alleles of the transcribed SNP were regulated differently across cell types (**Fig 4B**).

**Fig 4.**
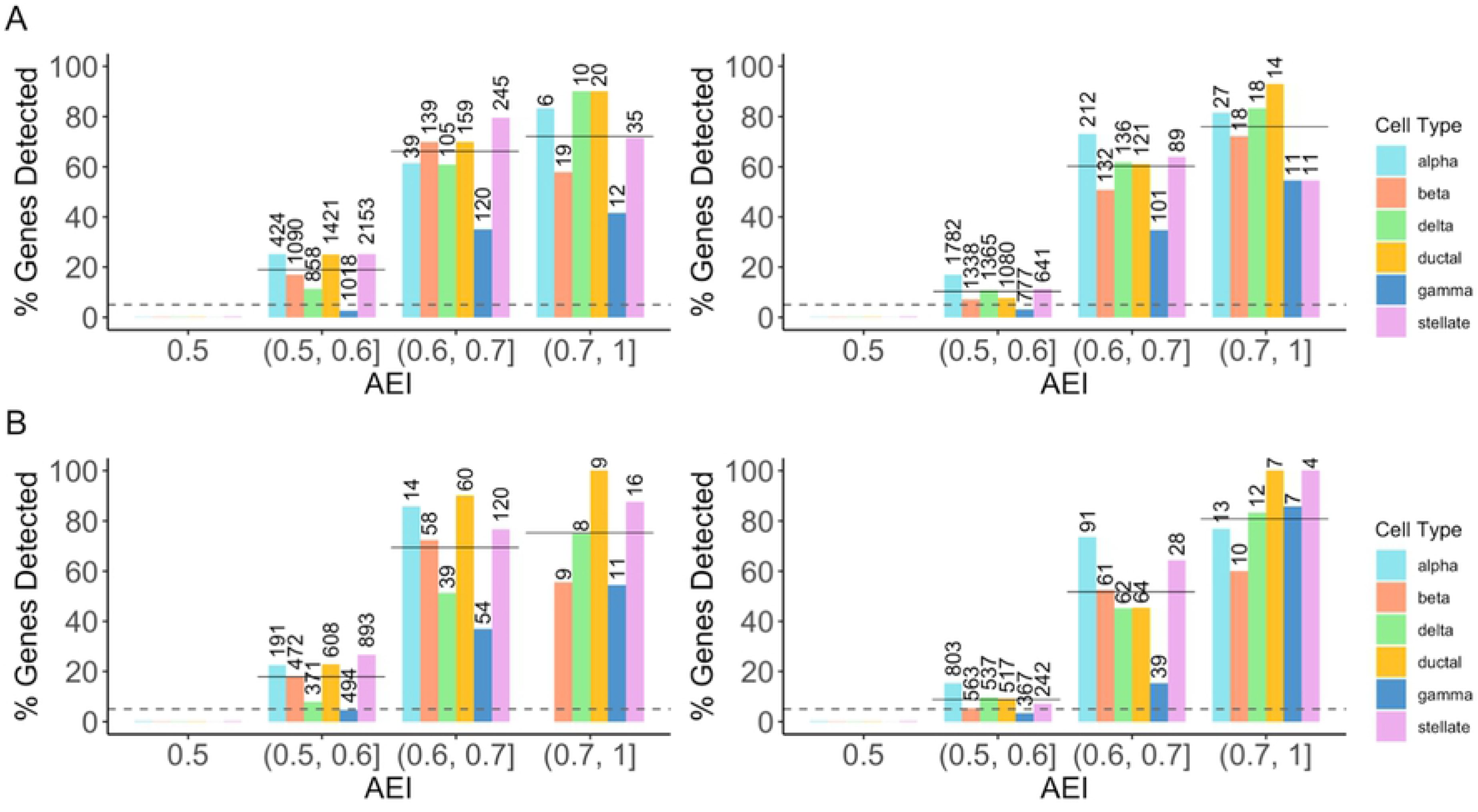
Benchmark evaluation for cell-type-specific AEI detection assuming two cell types with AEI. We evaluated the performance of BSCET assuming both the major and a non-major cell types had AEI using true cell type proportions. For each gene, the major cell type was selected as the one with the highest mean expression, and a non-major cell type was selected as the one with median mean expression in scRNA-seq data^9^. The AEI level for two cell types were assumed to be in the same direction, i.e., their AEI levels are > 0.5 for both cell types, for 70% of the SNPs, and in opposite directions, i.e., their AEI levels sum to 1, for the remaining 30% of the SNPs. **(A)** Type I error rate and power evaluated separately by cell type and true level of AEI for the 70% SNPs with the same direction of AEI under significance level *α* = 0.05 (dashed line) for the major **(left)** and non-major cell types **(right)**. **(B)** Type I error rate and power evaluated separately by cell type and true level of AEI for the 30% SNPs with opposite directions of AEI under significance level *α* = 0.05 (dashed line) for the major **(left)** and non-major cell types **(right)**. For each plot, the number above each bar indicates the total number of genes with cell-type-specific AEI within the category. The solid line indicates the overall power across all cell types.

In comparison, among the 7,873 SNPs with AEI in the same direction, the bulk GLM method correctly identified 4,840 SNPs with AEI, of which 2,609 overlapped with findings by BSCET. Since bulk-level AEI reflects the collective effect of AEI across all cell types, the bulk GLM method is expected to be more powerful when the goal is to detect the overall evidence of AEI. But still, BSCET was able to detect an additional 578 SNPs with cell-type-specific AEI for the major cell type and 422 SNPs for the non-major cell type that were missed by the bulk method. For the 3,427 SNPs with opposite AEI effects, the bulk GLM method detected 644 SNPs with AEI, and 444 of them overlapped with those detected by BSCET. This time, the bulk detection method was less powerful than BSCET, which was not surprising because the cell-type-specific AEI was neutralized when expression was averaged across cell types. For SNPs with bulk AEI level between 0.45 and 0.55, BSCET correctly uncovered many of them showing cell-type-specific AEI in both the major and non-major cell types, while the GLM model failed to detect any, indicating that BSCET was able to reveal cell-type-specific effect when the AEI was masked at the bulk level (**S3 Fig**). When repeating the same analysis but using the “estimated” cell type proportions as input for BSCET, for the 70% SNPs with the same AEI direction, we identified 1,766 SNPs for the major cell type and 1,119 for the non-major cell type after FDR adjustment, respectively. For the 30% SNPs with opposite AEI effects, BSCET correctly detected 707 for the major cell type and 375 for the non-major cell type. As expected, estimation uncertainty in cell type proportions slightly decreased BSCET’s detection power. But overall, the results were consistent with those using the true cell type proportions as input with the correlation of p-values over 0.9 (**S4 Fig**).

### Detection of cell-type-specific differential AEI

We further evaluated the performance of BSCET for detecting covariate effect on cell-type-specific AEI. We considered a case-control setting and assessed the effect of disease status on AEI, i.e., differential AEI (DAEI) between healthy and diseased samples. First, we considered equal number of cases and controls, i.e., non-diabetes and diabetes given the pancreatic islets data setting (100 vs. 100). **Fig 5A** shows that BSCET did not generate excessive false positive results for SNPs without cell-type-specific DAEI at the 0.05 significance level. When the difference of cell-type-specific AEI between cases and controls was small, e.g., 0.1, using true cell type proportions as input, among all 5,607 SNPs with cell-type-specific DAEI, BSCET identified 2,776 of them after multiple testing adjustment, whereas the bulk detection method uncovered 3,199 SNPs, with 2,136 overlapped with BSCET. When the differential AEI effect increased to 0.2, among the 5,655 SNPs with cell-type-specific DAEI, BSCET identified 4,370, while the bulk method identified 4,647, of which 3,981 overlapped with BSCET. Similar to one condition analysis, under both scenarios, SNPs uniquely detected by BSCET on average had lower sequencing depth as compared to SNPs uniquely detected by the bulk method. For example, when the cell-type-specific DAEI was 0.1, the mean read count for bulk-uniquely detected SNPs was 385.0 (IQR 133.4-479.8) and 98.4 (IQR 44.5-121.8) for BSCET-uniquely detected SNPs. The power decreased as the AEI level in controls increased. This was expected because given the same AEI difference between the two groups, a higher AEI level in controls led to a lower effect size of the AEI difference.

**Fig 5.**
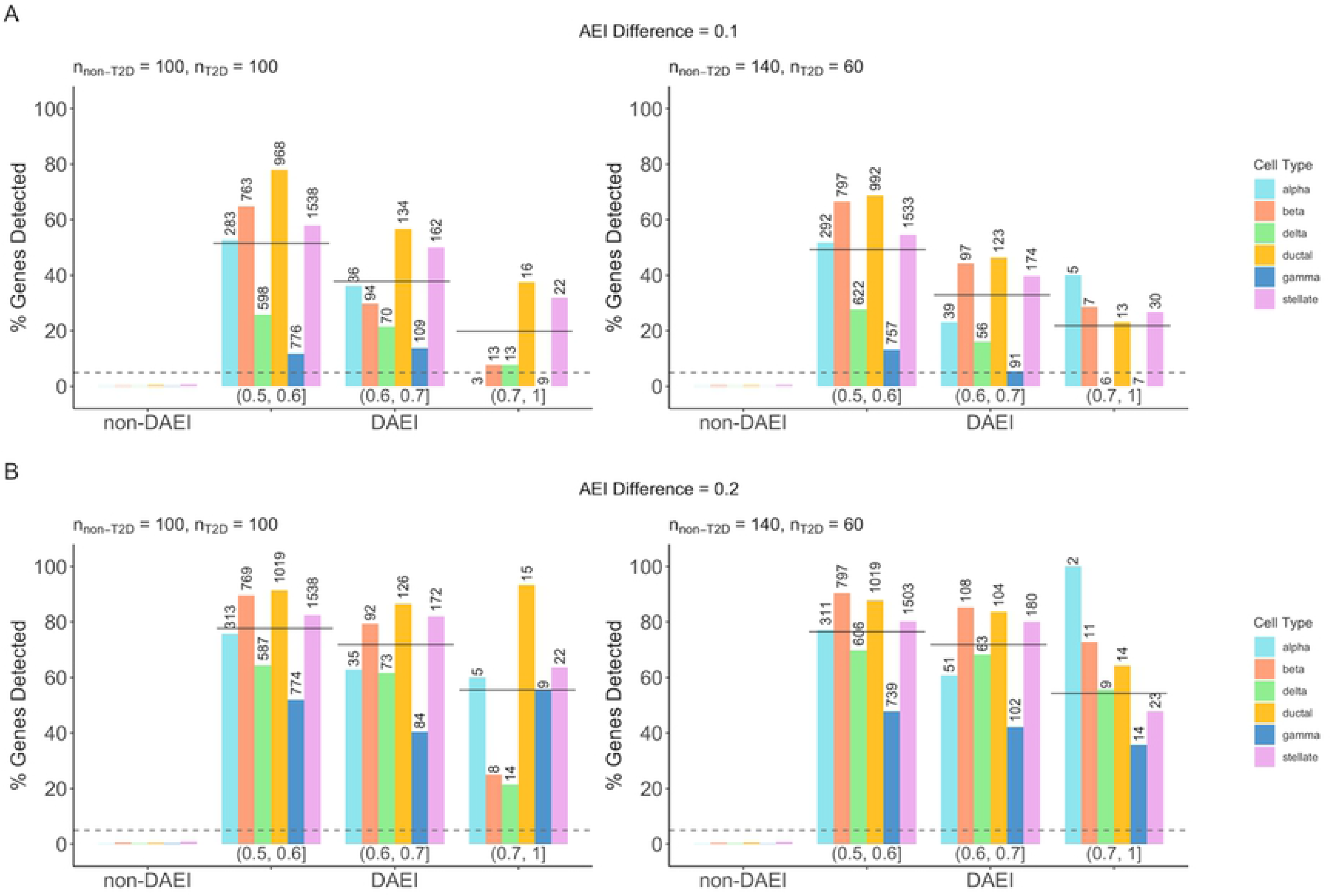
Benchmark evaluation for cell-type-specific differential AEI (DAEI) detection. Using true cell type proportions, we evaluated the performance of BSCET as a function of sample size for healthy (i.e., non-T2D) and diseased (i.e., T2D) samples, and true cell-type-specific AEI difference between two-condition samples (0.1 **(A)** and 0.2 **(B)**). Type I error rate (non-DAEI) and power (DAEI) were evaluated at significance level *α* = 0.05 (dashed line), further separated by the cell type and level of AEI in the healthy samples. The number above each bar indicates the total number of SNPs with cell-type-specific DAEI within the category. The solid line indicates the overall power across all cell types.

In real studies, we often have less cases than controls. Thus, we also considered a scenario assuming 60 cases versus 140 controls. Again, the type I error was under control. Compared to results with balanced sampling, this scenario had lower detection power, but BSCET still correctly detected 2,676 out of 5,641 SNPs with cell-type-specific DAEI when the true DAEI was 0.1, and 4,317 out of 5,656 when the true DAEI increased to 0.2 (**Fig 5B**). Additionally, when using “estimated” cell type proportions as input, across all scenarios, the results were highly correlated with those obtained using true cell type compositions (**S5 Fig**).

### Application to Fadista *et al.* human pancreatic islet bulk RNA-seq data

We applied BSCET to detect cell-type-specific AEI using human pancreatic islet bulk RNA-seq data of 89 donors generated in an expression quantitative trait loci (eQTL) study by Fadista *et al.*^12^. We chose to reanalyze this data because cell type compositions vary across individuals^11^, which may confound bulk-level AEI detection as one cannot tell whether the detected AEI is true AEI or is simply due to variations in cell type composition among individuals. We focused our analysis on four well-recognized cell types: acinar, alpha, beta, and ductal. We first deconvolved the bulk RNA-seq data by MuSiC^11^ using the scRNA-seq data from six healthy donors generated by Segerstolpe *et al.*^13^ as reference. The estimated cell type proportions were shown in **Fig 6A**. A SNP was included for analysis only if its minor allele count ≥ 5, total read count for both alleles ≥ 20, minor allele count was at least 5% of the total read count and appeared in at least 20 individuals. In total, we analyzed 5,972 SNPs across 2,909 genes using both BSCET and the bulk GLM method for AEI detection. After FDR adjustment, 283 SNPs across 129 genes were detected as having AEI by BSCET for at least one cell type (**S1 Table**). Of the 129 genes, 121 (94%) were identified while 8 (6%) were missed by the bulk method. The bulk method detected an additional 1,281 genes with AEI (**Fig 6B**). Similar to benchmark studies, the mean expression level was higher for SNPs detected only by bulk analysis (mean reads 129.3 with IQR 36.8-101.5) than SNPs detected only by BSCET (mean reads 46.7 with IQR 30.0-53.3).

**Fig 6.**
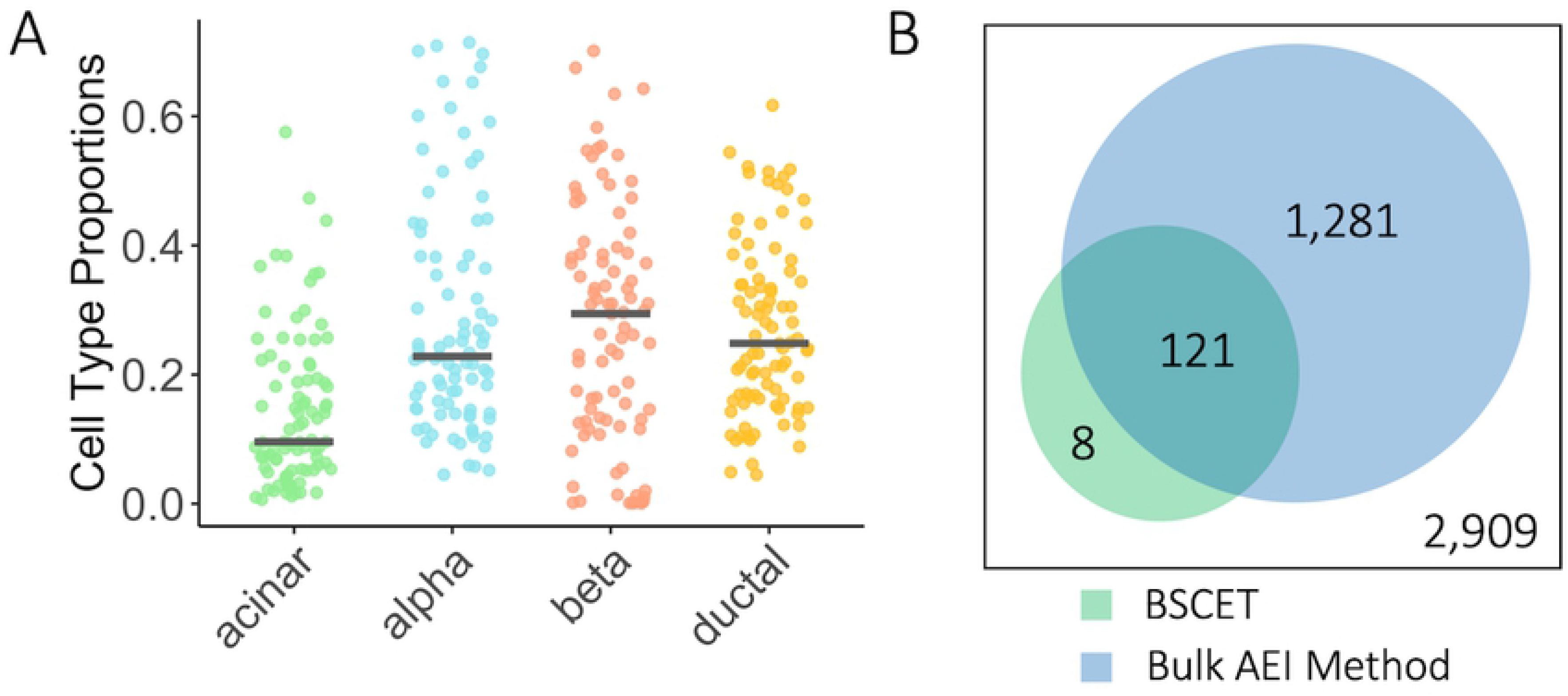
Deconvolution and cell-type-specific AEI detection of the Fadista pancreatic islets RNA-seq data. **(A)** Cell type proportion estimates of the Fadista bulk RNA-seq data^12^ using MuSiC^11^. The Segerstolpe scRNA-seq data^13^ were used as the reference. **(B)** Venn diagram showed the total number of genes analyzed, and number of genes detected with cell-type-specific AEI after FDR multiple testing adjustment using BSCET (green) and traditional bulk AEI detection method based on GLM model (blue).

Next, we compared genes detected by BSCET with the 616 eGenes identified to have *cis*-eQTL SNPs by Fadista *et al.*^12^. Among the 616 eGenes, 56 were analyzed by BSCET and 6 (11%) had cell-type-specific AEI. To evaluate if 11% overlapping was higher than expected by chance, we performed resampling-based enrichment analysis. Specifically, we randomly sampled 129 genes from the remaining 2,780 genes that did not show evidence of cell-type-specific AEI by BSCET and recorded the percentage of genes overlapping with the 616 eGenes in Fadista *et al.*^12^. By repeating this sampling procedure 10,000 times, we calculated the enrichment p-value as 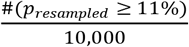, which equaled to 0.029, suggesting that results from BSCET significantly overlapped with the *cis*-eQTL findings.

For genes identified with cell-type-specific AEI by BSCET, some of them are cell-type-specific marker genes by PanglaoDB^14^. For example, *CELA3A*, with two transcribed SNPs detected having cell-type-specific AEI by BSCET, is a marker gene for acinar cells (**Fig 7A**), and BSCET only detected AEI in acinar cells but not in the other cell types, confirming the specificity of BSCET. Furthermore, we uncovered two other marker genes for acinar cells with cell-type-specific AEI, *CPA2* and *PRSS3* (**S6 Fig**). In addition to acinar cells, we also detected a marker gene for beta cells with cell-type-specific AEI, *EDARADD*, which had consistent signals across the majority of transcribed SNPs in this gene (**Fig 7B**).

**Fig 7.**
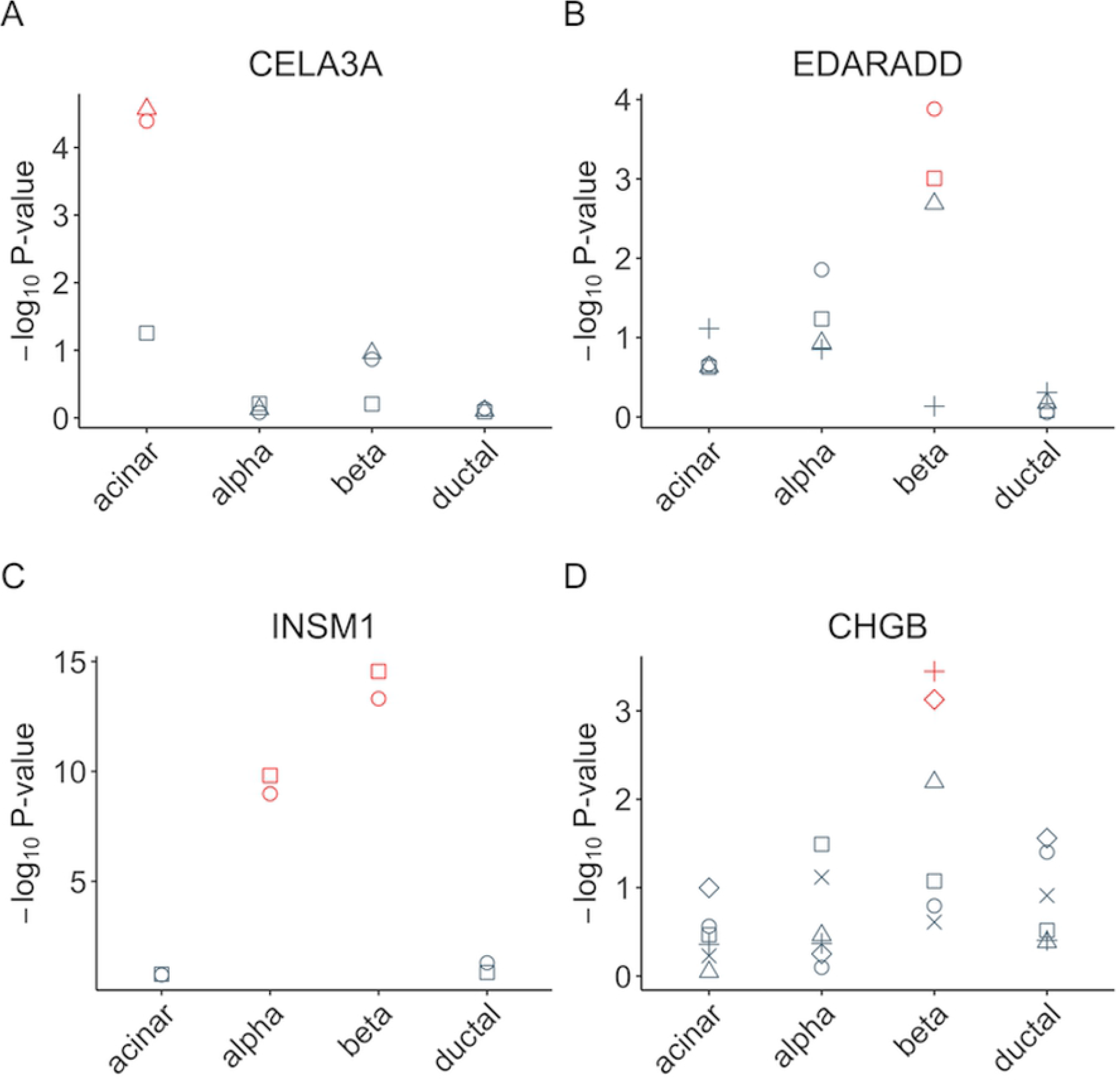
Selected genes with cell-type-specific AEI of the Fadista pancreatic islets RNA-seq data. We selected 4 genes, *CELA3A* **(A)**, *EDARADD* **(B)**, *INSM1* **(C)** and *CHGB* **(D)**, to show their SNP-level p-values of cell-type-specific AEI. Within each cell type, different shapes represent different SNPs of the gene, with red color indicating significant AEI after FDR multiple testing adjustment.

For genes identified by BSCET but are not documented as marker genes by PanglaoDB^14^, we found many of them are biologically relevant. For example, *INSM1* had cell-type-specific AEI for both alpha and beta cells with signals well agreed between the two SNPs analyzed for this gene (**Fig 7C**). Several studies have reported *INSM1* as a key factor for the development and differentiation in pancreatic beta cells^15, 16^. Another example is *CHGB*, which had significant cell-type-specific AEI for beta cells by BSCET (**Fig 7D**). A recent study has shown that *CHGB* is essential for regulating secretory granule trafficking in beta cells^17^. Several other genes identified by BSCET are also known to have relevant functions. For example, *PPM1K* elevates branched-chain amino acids concentrations and is related to high risk of T2D^18, 19^. *SYT11* has been reported to be positively related to glucose stimulated insulin secretion in human pancreatic islets^20^. *ERO1B* is known to encode a pancreas-specific disulfide oxidase that promotes protein folding in pancreatic beta cells^21^. *AHR* is expressed in aortic endothelial cells and is activated in response to high glucose stimulation^22^ (**S6 Fig**).

### Evaluation on the impact of scRNA-seq reference

To examine whether BSCET is robust to the input cell type compositions, we reanalyzed the Fadista *et al.* bulk RNA-seq data using a different scRNA-seq dataset as reference. The new scRNA-seq dataset was generated from three healthy donors by Baron *et al.*^9^. Cell type deconvolution using this scRNA-seq reference resulted in a much smaller proportion for beta cells compared to results obtained using the Segerstolpe scRNA-seq data^13^ as reference (**S7A Fig**). Using the new cell type proportions as input for BSCET, we identified 148 out of 2,909 analyzed genes to have cell-type-specific AEI, among which 102 were also found in previous analysis when using the Segerstople scRNA-seq data^13^ as reference (**S2 Table**). Overall, the SNP level p-values obtained from these two different scRNA-seq reference datasets were well correlated (R = 0.72, **S7B Fig**). Stratified by cell type, the correlation was the highest for alpha cells (R = 0.87), followed by acinar (R = 0.81) and ductal (R = 0.77) cells. The correlation for beta cells was moderate (R = 0.59), which can be explained by the large difference in the estimated beta cell proportions when using the two single-cell datasets as reference (mean proportion: 3% vs. 28%, **S7C Fig**). Overall, BSCET yielded relatively consistent results, suggesting that it is robust to the choice of single-cell reference for deconvolution.

### Replication using Bunt *et al.* human pancreatic islet bulk RNA-seq data

We further applied BSCET to another pancreas islet RNA-seq data of an eQTL study generated from 118 individuals by Bunt *et al.*^23^ using the Segerstolpe scRNA-seq data^13^ as reference. The estimated cell type compositions are overall similar to the Fadista data^12^, except for a smaller proportion for alpha cells (**S8 Fig**). In total, we analyzed 9,013 SNPs across 4,293 genes, and detected 722 SNPs across 423 genes showing cell-type-specific AEI after multiple testing adjustment (**S3 Table**). Next, we compared these results with those obtained from the Fadista data^12^. Among the 48 genes detected with AEI for acinar cells in Fadista, 41 were analyzed in Bunt, among which 23 (56%) also had AEI in acinar cells. Similarly, for the 81 genes with AEI for beta cells in Fadista, 33 (41%) were replicated in Bunt. For ductal cells, among the 27 genes with AEI in Fadista, 15 (56%) were replicated in Bunt. However, for alpha cells, among the 39 genes detected in Fadista, only 6 genes (15%) were replicated in Bunt. This is likely due to the large discrepancy in estimated cell type proportions for alpha cells between the two datasets. Despite some difference, the AEI results obtained using the Bunt data validated findings obtained in Fadista. For example, *PVR*, also known as *CD155*, involves in the regulation of T-cell activation and is associated with autoimmune diseases such as type 1 diabetes (T1D)^24, 25^, had two SNPs showing AEI for ductal cells in Fadista. This finding was confirmed in Bunt in multiple SNPs (**Fig 8A**). Similarly, we detected acinar and beta cell-specific AEI for several SNPs in gene *SSR3* in both datasets (**Fig 8B**). Gene *SSR3* is involved in protein translocation across the endoplasmic reticulum (ER) membrane, whose expression has showed to be related to T2D^26, 27^. Moreover, gene *GNPNAT1* was detected only for one SNP with acinar cell-specific AEI in Fadista, but was confirmed with much stronger effect in Bunt (**Fig 8C**). Previous studies have shown that *GNPNAT1 is* associated with insulin secretion and diabetes^28, 29^.

**Fig 8.**
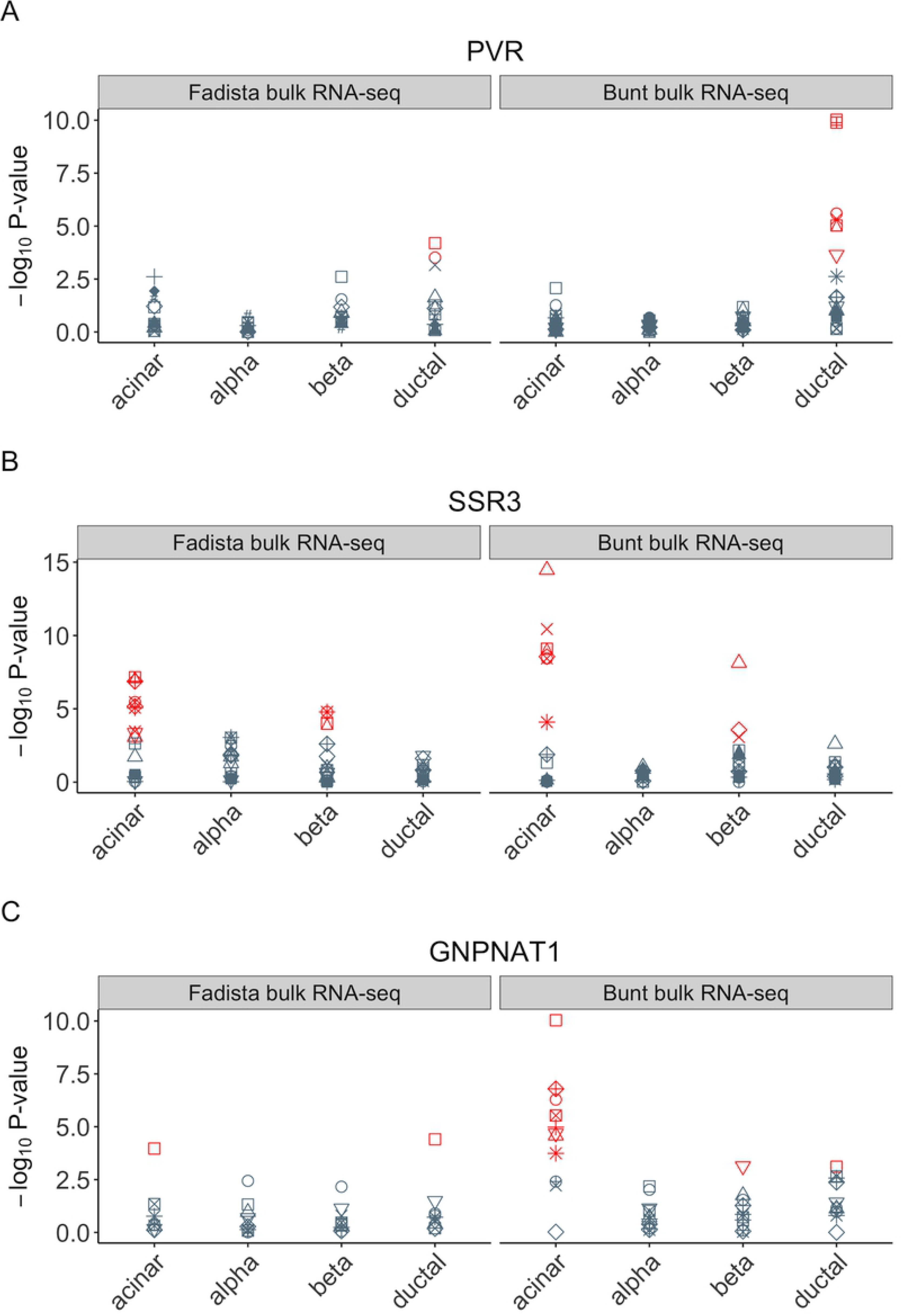
Selected genes with cell-type-specific AEI of both pancreatic islet RNA-seq data. We selected 3 genes, *PVR* **(A)**, *SSR3* **(B)** and *GNPNAT1* **(C)**, to show their SNP-level p-values of cell-type-specific AEI using two different bulk samples, Fadista *et al.*^12^ and Bunt *et al.*^23^. Within each cell type, different shapes represent different SNPs of the gene, with red color indicating significant AEI after FDR multiple testing adjustment.

### Association of cell-type-specific AEI with HbA1c level

Type 2 diabetes (T2D) is characterized by the progressive dysfunction of pancreas islets. Next, we applied BSCET to explore the association between hemoglobin A1c (HbA1c), a well-known biomarker for T2D diagnosis, and cell-type-specific AEI. People with higher HbA1c have greater risk of developing T2D^30^. Using the Fadista bulk RNA-seq data, we focused our analysis on the 77 samples with HbA1c level available. **Fig 9A** showed relationship between HbA1c level and estimated cell type proportions with Segerstolpe scRNA-seq data^13^ as reference. As expected, we observed a negative association between beta cell proportion and the HbA1c level. After applying the same filtering criteria as described previously, we analyzed a total of 5,021 SNPs across 2,570 genes, and identified 8 genes, *HYOU1*, *PLA2G1B*, *CCDC32*, *CCL2*, *CDC42EP3*, *LARS*, *SLC30A8* and *CEL*, whose cell-type-specific AEI was associated with HbA1c level (**S4 Table**). By contrast, using the bulk AEI detection method, we detected 125 genes with significant association between AEI and HbA1C level, and among which, *LARS*, *SLC30A8* and *CEL* overlapped with those detected by BSCET.

**Fig 9.**
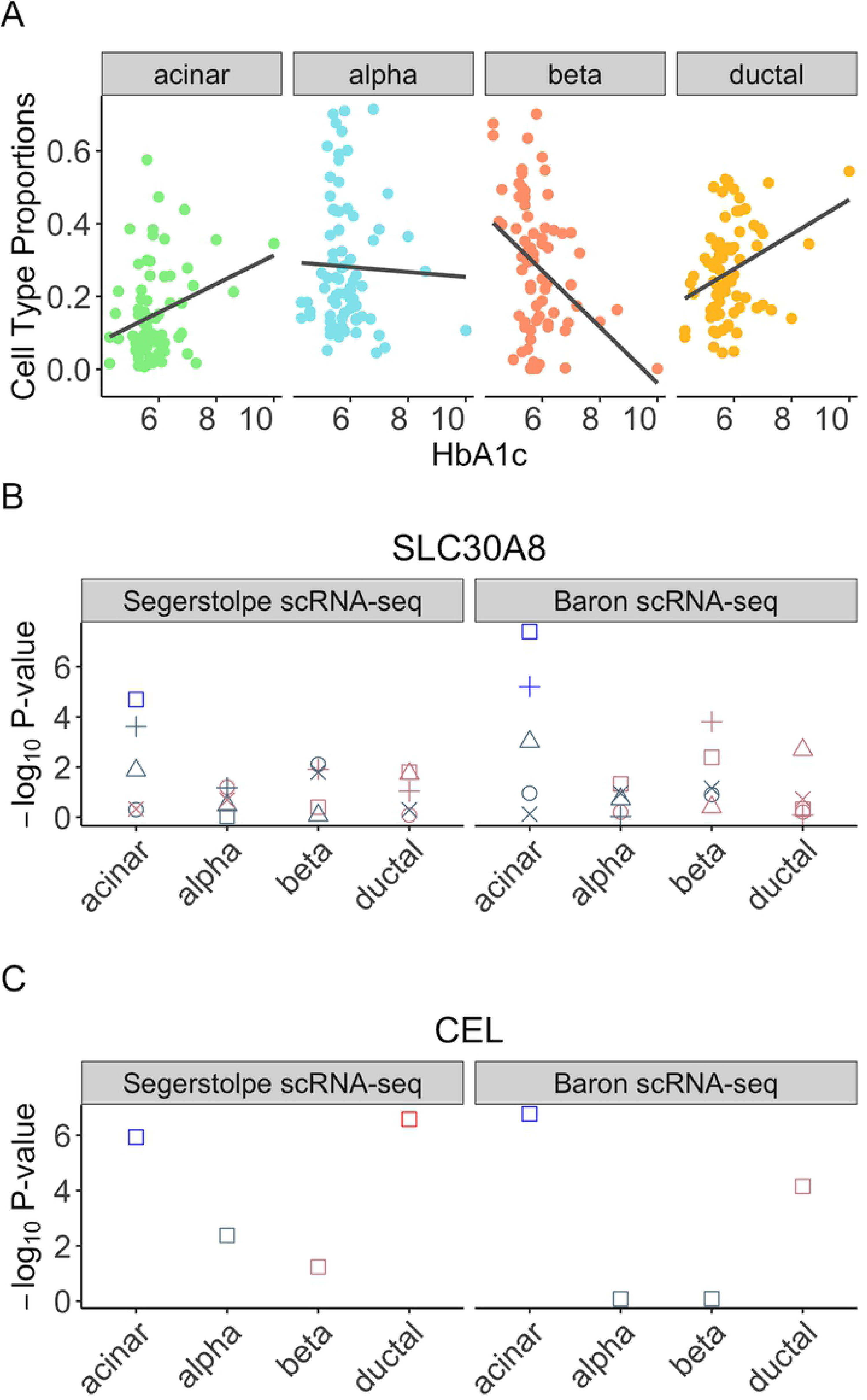
Cell type compositions and selected genes with cell-type-specific AEI associated with the progression of T2D of the Fadiata pancreatic islets RNA-seq data. **(A)** HbA1c vs. estimated proportions of the Fadista bulk RNA-seq data^12^ for each cell type, where the proportions were estimated through MuSiC^11^ based on the Segerstolpe single-cell reference^13^. **(B, C)** We selected 2 genes, *SLC30A8* **(B)** and *CEL* **(C)**, to show their SNP-level p-values of the association between HbA1c and cell-type-specific AEI, where we used cell type proportion estimates obtained from two different scRNA-seq reference datasets: Segerstolpe^13^ and Baron^9^. Within each cell type, different shapes represent different SNPs. Red color indicates a positive correlation between HbA1c and cell-type-specific AEI, with color brightness representing significance level. Similarly, blue color blue indicates a negative correlation between HbA1c and cell-type-specific AEI, with color brightness representing significance level.

BSCET revealed a negative association of HbA1c level with the acinar cell-specific AEI for *SLC30A8* (p-value = 0.00002) (**Fig 9B**). This is consistent with previous studies reporting that *SLC30A8* encodes an islet zinc transporter (ZnT8) and a reduced zinc transport can increase the risk of T2D^31, 32^. Moreover, a human study suggested that *SLC30A8* haploinsufficiency helps protect against T2D^33^. To further validate our findings, we repeated the analysis but using the cell type composition estimated with the Baron scRNA-seq data^9^ as reference (**S9A Fig**). After multiple testing adjustment, we identified 4 genes, *CCDC32*, *LARS*, *SLC30A8* and *CEL*, whose cell-type-specific AEI was associated with HbA1c level, and all were also detected when using the Segerstolpe data as single-cell reference (**S5 Table**). For *SLA30A8*, we observed a consistent negative association between acinar cell-specific AEI and HbA1c level. Similar negative association of acinar cell-specific AEI was detected for another gene, *CEL* (**Fig 9C**), a marker gene for acinar cells in PanglaoDB^14^. Previous studies have reported that *CEL* mutations can lead to childhood-onset pancreatic exocrine dysfunction and diabetes mellitus from adulthood^34, 35^. For the other two genes, *CCDC32* and *LARS*, further investigations are needed as little is known about their functions related to T2D (**S9 Fig**).

## Discussion

Detection of AEI is an important step towards the understanding of phenotypic variations associated with gene expression differences across individuals. Traditionally, AEI is detected at the tissue level using bulk RNA-seq data. However, solid tissue includes cells from different cell types and gene expression variations are often cell-type-specific. Methods that detect AEI at the bulk tissue level only capture the averaged effect of gene expression across cells, making it difficult to discern what cell types are driving the AEI signal. Although it is possible to study cell-type-specific AEI using scRNA-seq data, the high cost of scRNA-seq has limited its application in studies that involve a large number of individuals. As bulk RNA-seq is more cost-effective and bulk RNA-seq data in clinically relevant tissues are widely available, to better utilize existing bulk RNA-seq data and help refine bulk-level AEI, we developed BSCET, a novel method that can detect cell-type-specific AEI across individuals. BSCET is not aimed to replace the traditional bulk AEI detection methods, but rather is set to help elucidate in what cell types AEI is present. To gain a comprehensive understanding of genetic polymorphisms on gene expression variation, we suggest analyzing the data using both the traditional bulk tissue AEI detection methods and BSCET.

To detect cell-type-specific AEI at the population level, BSCET aggregates information across individuals of the same transcribed SNP. A major challenge for cross-individual AEI detection is the difficulty of aligning allele-specific read counts, as the corresponding regulatory SNP for a given gene is often not observed. In BSCET, we assumed the unobserved regulatory SNP is in complete linkage disequilibrium with the transcribed SNP, which allows us to align read counts from different individuals based on alleles of the transcribed SNP. The aligned allelic read counts enabled the detection of AEI across individuals at the population level. Through extensive benchmark evaluations, we showed that BSCET had adequate power to detect moderate AEI at the cell type level. Through further applications to human pancreatic islet RNA-seq datasets, we demonstrated that genes with cell-type-specific AEI uncovered by BSCET are biologically relevant to pancreatic functions and the progression of T2D.

BSCET uses estimated cell type proportions of bulk RNA-seq data as input. Ideally, if the true cell type proportions are known, we would like to use them in order to obtain an accurate estimate of cell-type-specific AEI. In the absence of true cell type proportions, we can estimate them using cell type deconvolution methods such as MuSiC^11^, which borrows information from external scRNA-seq data to infer the cell type composition of a bulk RNA-seq sample. The benchmark evaluations and real data applications showed that the results obtained using “estimated” cell type proportions were highly correlated with those obtained using true proportions. We further illustrated that, even though different scRNA-seq reference can lead to varied proportion estimates, such variations did not affect the BSCET results much as the detected AEI genes agreed well, suggesting that BSCET is robust to estimation uncertainty on the input.

As a regression-based method, BSCET is flexible and can include covariates in the model and evaluate how covariates would affect cell-type-specific AEI. We have demonstrated BSCET has adequate power to detect association between cell-type-specific AEI with covariates through benchmark evaluations under a variety of scenarios. Further, we applied the extension model to human pancreatic islet bulk RNA-seq data and detected genes whose cell-type-specific AEI is associated with T2D progression. We note that as more parameters are included in the model, a larger sample size is required. Therefore, we would recommend users to perform such analysis as a supplemental step only after evidence of cell-type-specific AEI is detected in at least one of the two groups or in all individuals in the data.

In summary, we have developed BSCET, a regression-based method that integrates bulk RNA-seq and scRNA-seq data to detect cell-type-specific AEI across individuals in a population. BSCET refines the current bulk-level AEI detection workflow and helps understand gene regulation and its association with phenotypic variations across individuals at cell type resolution. As bulk RNA-seq is widely adopted by many biomedical studies and more samples are being sequenced, instead of generating large samples using scRNA-seq, it is desirable for researchers to fully utilize the existing bulk RNA-seq data and learn a comprehensive picture of allelic imbalance in a more cost-effective way. We believe BSCET, which, to the best of our knowledge, is the first method using bulk RNA-seq data to detect cell-type-specific AEI, will be a great supplemental tool for utilizing the easily accessible bulk sequencing data to elucidate cell type contributions in human diseases.

## Methods

BSCET takes two datasets as input: 1) a bulk RNA-seq dataset in which bulk level gene expression is measured in a relatively large set of individuals; and 2) a scRNA-seq dataset with cells generated from the same tissue as the bulk RNA-seq data in a small set of individuals. The scRNA-seq dataset can be from the same individuals in the bulk RNA-seq dataset, or from different individuals obtained from an external dataset. The goal of BSCET is to detect cell-type-specific AEI in the bulk RNA-seq samples. BSCET involves two steps, and an overview of the procedure is shown in **Fig 1**.

### Step 1: Estimating cell type proportions by deconvolution

Since cell type proportions are often unknown in bulk RNA-seq samples, in this step, we aim to infer cell type proportions in the bulk RNA-seq samples by incorporating cell-type-specific gene expression information provided by a scRNA-seq reference. This can be achieved by cell type deconvolution algorithms such as MuSiC^11^. Following MuSiC^11^, for gene *g* of individual *j*, the total bulk-level read count, *X*_*jg*_, can be written as a weighted sum of *K* cell-type-specific gene expressions,

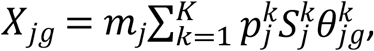

where, for individual *j*, *m_j_* is the total number of cells in the tissue for bulk RNA-seq, 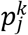 is the proportion of cells from cell type *k*, 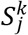 is the average cell size of cell type *k*, and 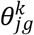 is the relative abundance of gene *g* for cell type *k*. Since subject-level 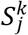 and 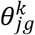 are often not available, by borrowing information from external scRNA-seq reference, they can be approximated by the sample mean, *S^k^* and 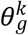. For individual *j*, as *m_j_* is a constant across all genes, the cell type proportions, 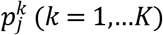, can be inferred through a weighted non-negative least squares regression by regressing the bulk-level expression across all genes, *X*_*jg*_ (*g* = 1,…,*G*), on the cell-type-specific gene expression, 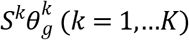. MuSiC^11^ has shown superior performance in estimating cell type proportions than popular methods such as CIBERSORT^36^.

### Step 2: Detecting cell-type-specific AEI

To detect cell-type-specific AEI across individuals in a population, we consider one transcribed SNP at a time. For ease of notation, we omit index for genes. Let *T* and *t* be the two alleles at a transcribed SNP. For individual *j*, let 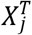 and 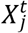 be the read counts for the *T* and *t* alleles, respectively. Following MuSiC^11^, the bulk-level read count of each transcribed allele can be written as a weighted sum of the *K* cell type-specific allelic expressions,

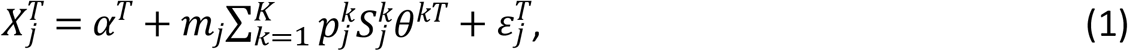

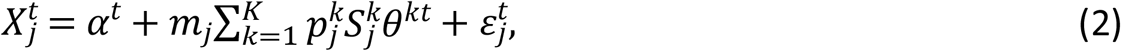

where *α*^*T*^ and *α*^*t*^ are intercepts to capture the information not explained by the *K* cell types, 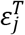 and 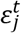 are independent random errors assumed to follow *N*(0,*σ*^2^), and *θ*^*kT*^ and *θ*^*kt*^ are mean expression of the transcribed alleles, for cell type *k*, across individuals in the population. Taking difference between (1) and (2), we get

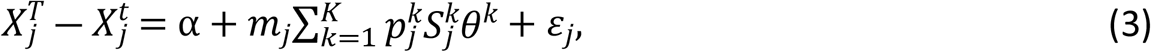

where *α* = *α*^*T*^ − *α*^*t*^ represents allelic expression difference not explained by the selected *K* cell types, and *θ^k^* = *θ*^*kT*^ − *θ*^*kt*^ represents allelic expression difference between alleles *T* and *t* in cell type *k*.

We expect the two transcribed alleles to be equally expressed, i.e., *θ^k^* = 0, in the absence of cell-type-specific AEI, and *θ^k^* ≠ 0 otherwise. Therefore, to detect cell-type-specific AEI in the population, for cell type *k*, we test the following hypothesis,

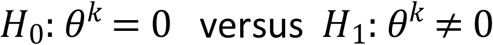

using a *t* statistic.

Although we can use estimated cell type proportions obtained from MuSiC^11^ to approximate 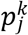, *m_j_* and 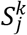 are not directly observed. To circumvent this, we use the individual-specific library size factor obtained from DESeq2^37^ as an estimate for *m_j_*. For 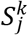, we borrow the idea from MuSiC^11^ and approximate it by *S^k^*, assuming that cell size of all individuals is equal for a given cell type. Since *S^k^* is a constant across all individuals, it is considered as a nuisance parameter and not estimated.

### Step 2 extension: Association between cell-type-specific AEI and covariates

As a regression-based method, we can readily extend it to assess covariate effects on cell-type-specific AEI. For illustration purposes, we only consider one covariate, but it is straightforward to incorporate multiple covariates. Let *V_j_* be the covariate of interest for individual *j*. We modify model (3) by adding an interaction term between the covariate and the estimated cell type proportions:

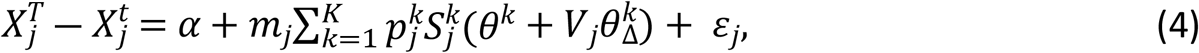

where *θ^k^* is the population-level AEI of the transcribed SNP for cell type *k* controlling for the covariate; 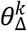 is the covariate effect on the cell-type-specific AEI, interpreted as the change in population-level AEI for cell type *k* as the covariate increases by 1 unit; *ε*_*j*_ is the random error term and assumed to follow *N*(0,*σ*^2^). For example, in a case-control study where the covariate indicates disease status (e.g., coded as 1 for cases), *θ^k^* is the cell-type-specific AEI in controls, and 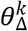 is the difference in the cell-type-specific AEI between cases and controls, a.k.a. cell-type-specific differential AEI.

In practice we will likely test for the covariate effect on cell-type-specific AEI only if a cell-type-specific AEI has been detected based on model (3). Therefore, in model (4), we are interested in testing whether the cell-type-specific AEI changes with the covariate, i.e., for each cell type *k*, we test the following hypothesis,

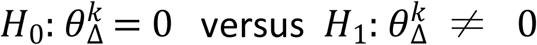

using a *t* statistic.

### Benchmark bulk RNA-seq data generation

To evaluate the performance of BSCET, we constructed benchmark bulk RNA-seq data in which the cell type proportions and AEI levels for each cell type are known. The benchmark data include 60 individuals generated using a publicly available scRNA-seq dataset on human pancreatic islets from three healthy individuals^9^. For each individual in the benchmark bulk RNA-seq data, we generated 5,000 single cells from six selected cell types, including alpha, beta, delta, ductal, gamma and stellate. The cell type proportions were assumed to follow a Dirichlet distribution, with mean proportions inferred from the original scRNA-seq data^9^ (**Fig 2A**).

Since BSCET analyzes one transcribed SNP at a time, for the benchmark bulk RNA-seq data, we assumed each gene only has one transcribed SNP. For each cell, we simulated read counts for 11,300 SNPs, corresponding to the number of genes expressed in the original scRNA-seq data. To better reflect patterns seen in real data, for each cell type, we generated read counts only for a fraction of cells, with the percentage of expressed cells inferred from the original scRNA-seq data. **Fig 2B** shows the overall workflow for benchmark dataset generation. Briefly, for each gene, we obtained the mean cell-type-specific expression across cells from the original scRNA-seq data. This mean value set the Poisson distribution rate parameter and subject-specific mean expression was sampled from this Poisson distribution. To generate cell-specific total read count for each individual, we sampled from another layer of Poisson distribution where the rate parameter was determined from the previous step. The total read count of each transcribed SNP was then split into two allele-specific read counts through a binomial distribution with the probability set by the predefined cell-type-specific AEI for the gene. For cell types without AEI, we set their AEI level to 0.5 for all genes. For cell types with AEI, we assumed their AEI levels for each gene followed a truncated gamma distribution with the majority of genes having small to moderate AEI. By summing up allelic read counts across all cells, we obtained the benchmark bulk-level gene expression data.

We also generated benchmark data that include samples from two conditions (e.g. healthy versus diseased). This allows us to evaluate the performance of BSCET in detecting covariate effect on cell-type-specific AEI. Using similar data generation scheme as described above, we generated artificial single-cell and bulk RNA-seq data for 200 individuals, with the ratio of healthy and diseased being 1:1 or 7:3. For each gene, we assumed only the major cell type, i.e., the cell type with the highest mean expression among all cell types, had AEI, and 50% of the genes had differential AEI between healthy and diseased individuals. For genes with cell-type-specific differential AEI, the AEI level for diseased individuals was assumed to be higher than that of healthy individuals by ∆_*AEI*_, which took two possible values, 0.1 and 0.2.

## Acknowledgements

This work was supported by the following grants: R01GM125301 (to M.L.), R01HL113147 (to M.L.), R01HL150359 (to M.L.), and R01EY030192 (to M.L.).

## Web Resources

A tutorial of BSCET is available at https://jiaxin-fan.github.io/BSCET.github.io/Introduction.html.

## References

1. Pastinen, T., Hudson, T.J. (2004). Cis-Acting Regulatory Variation in the Human Genome. Science 306, 647.

2. Sun, W., Hu, Y. (2013). eQTL Mapping Using RNA-seq Data. Stat Biosci 5, 198–219.

3. Edsgärd, D., Iglesias, M.J., Reilly, S., Hamsten, A., Tornvall, P., Odeberg, J., Emanuelsson, O. (2016). GeneiASE: Detection of condition-dependent and static allele-specific expression from RNA-seq data without haplotype information. Scientific reports 6, 21134.

4. Mayba, O., Gilbert, H.N., Liu, J., Haverty, P.M., Jhunjhunwala, S., Jiang, Z., Watanabe, C., Zhang, Z. (2014). MBASED: allele-specific expression detection in cancer tissues and cell lines. Genome Biol. 15, 405.

5. Fan, J., Hu, J., Xue, C., Zhang, H., Susztak, K., Reilly, M.P., Xiao, R., Li, M. (2020). ASEP: Gene-based detection of allele-specific expression across individuals in a population by RNA sequencing. PLOS Genetics 16, e1008786.

6. Tibshirani, R., Perry, N.M., Khatri, P., Butte, A.J., Davis, M.M., Bodian, D.L., Staedtler, F., Hastie, T., Shen-Orr, S.S., Sarwal, M.M. (2010). Cell type-specific gene expression differences in complex tissues. Nature Methods 7, 287–289.

7. Handley, A., Schauer, T., Ladurner, A.G., Margulies, C.E. (2015). Designing Cell-Type-Specific Genome-wide Experiments. Mol. Cell 58, 621–631.

8. Hwang, B., Lee, J.H., Bang, D. (2018). Single-cell RNA sequencing technologies and bioinformatics pipelines. Exp. Mol. Med. 50, 96.

9. Baron, M., Veres, A., Wolock, S., Faust, A., Gaujoux, R., Vetere, A., Ryu, J., Wagner, B., Shen-Orr, S., Klein, A. et al. (2016). A Single-Cell Transcriptomic Map of the Human and Mouse Pancreas Reveals Inter- and Intra-cell Population Structure. Cell Systems 3, 346–360.e4.

10. Lawlor, N., George, J., Bolisetty, M., Kursawe, R., Sun, L., Sivakamasundari, V., Kycia, I., Robson, P., Stitzel, M.L. (2017). Single-cell transcriptomes identify human islet cell signatures and reveal cell-type-specific expression changes in type 2 diabetes. Genome Res. 27, 208–222.

11. Wang, X., Park, J., Susztak, K., Zhang, N.R., Li, M. (2019). Bulk tissue cell type deconvolution with multi-subject single-cell expression reference. Nature Communications 10, 380.

12. Fadista, J., Vikman, P., Laakso, E.O., Mollet, I.G., Esguerra, J.L., Taneera, J., Storm, P., Osmark, P., Ladenvall, C., Prasad, R.B. et al. (2014). Global genomic and transcriptomic analysis of human pancreatic islets reveals novel genes influencing glucose metabolism. Proc. Natl. Acad. Sci. USA 111, 13924.

13. Segerstolpe, Å, Palasantza, A., Eliasson, P., Andersson, E., Andréasson, A., Sun, X., Picelli, S., Sabirsh, A., Clausen, M., Bjursell, M.K. et al. (2016). Single-Cell Transcriptome Profiling of Human Pancreatic Islets in Health and Type 2 Diabetes. Cell Metabolism 24, 593–607.

14. Franzén, O., Gan, L., Björkegren, J., L.M. (2019). PanglaoDB: a web server for exploration of mouse and human single-cell RNA sequencing data. Database (Oxford) 2019.

15. Gierl, M.S., Karoulias, N., Wende, H., Strehle, M., Birchmeier, C. (2006). The zinc-finger factor Insm1 (IA-1) is essential for the development of pancreatic beta cells and intestinal endocrine cells. Genes Dev. 20, 2465–2478.

16. Zhang, T., Wang, H., Saunee, N.A., Breslin, M.B., Lan, M.S. (2010). Insulinoma-associated antigen-1 zinc-finger transcription factor promotes pancreatic duct cell trans-differentiation. Endocrinology 151, 2030–2039.

17. Bearrows, S.C., Bauchle, C.J., Becker, M., Haldeman, J.M., Swaminathan, S., Stephens, S.B. (2019). Chromogranin B regulates early-stage insulin granule trafficking from the Golgi in pancreatic islet β-cells. J. Cell. Sci. 132, jcs231373.

18. Goni, L., Qi, L., Cuervo, M., Milagro, F.I., Saris, W.H., MacDonald, I.A., Langin, D., Astrup, A., Arner, P., Oppert, J. et al. (2017). Effect of the interaction between diet composition and the PPM1K genetic variant on insulin resistance and β cell function markers during weight loss: results from the Nutrient Gene Interactions in Human Obesity: implications for dietary guidelines (NUGENOB) randomized trial. ajcn 106, 902–908.

19. Lotta, L.A., Scott, R.A., Sharp, S.J., Burgess, S., Luan, J., Tillin, T., Schmidt, A.F., Imamura, F., Stewart, I.D., Perry, J.R.B. et al. (2016). Genetic Predisposition to an Impaired Metabolism of the Branched-Chain Amino Acids and Risk of Type 2 Diabetes: A Mendelian Randomisation Analysis. PLOS Medicine 13, e1002179.

20. Andersson, S.A., Olsson, A.H., Esguerra, J.L.S., Heimann, E., Ladenvall, C., Edlund, A., Salehi, A., Taneera, J., Degerman, E., Groop, L. et al. (2012). Reduced insulin secretion correlates with decreased expression of exocytotic genes in pancreatic islets from patients with type 2 diabetes. Mol. Cell. Endocrinol. 364, 36–45.

21. Awazawa, M., Futami, T., Sakada, M., Kaneko, K., Ohsugi, M., Nakaya, K., Terai, A., Suzuki, R., Koike, M., Uchiyama, Y. et al. (2014). Deregulation of pancreas-specific oxidoreductin ERO1β in the pathogenesis of diabetes mellitus. Mol. Cell. Biol. 34, 1290–1299.

22. Dabir, P., Marinic, T.E., Krukovets, I., Stenina, O.I. (2008). Aryl hydrocarbon receptor is activated by glucose and regulates the thrombospondin-1 gene promoter in endothelial cells. Circ. Res. 102, 1558–1565.

23. van de Bunt, M., Manning Fox, J.E., Dai, X., Barrett, A., Grey, C., Li, L., Bennett, A.J., Johnson, P.R., Rajotte, R.V., Gaulton, K.J. et al. (2015). Transcript Expression Data from Human Islets Links Regulatory Signals from Genome-Wide Association Studies for Type 2 Diabetes and Glycemic Traits to Their Downstream Effectors. PLOS Genetics 11, e1005694.

24. Lozano, E., Joller, N., Cao, Y., Kuchroo, V.K., Hafler, D.A. (2013). The CD226/CD155 interaction regulates the proinflammatory (Th1/Th17)/anti-inflammatory (Th2) balance in humans. Journal of immunology (Baltimore, Md.: 1950) 191, 3673–3680.

25. Escalante Nichole, K., von, R.A., Martin, L., Choy Jonathan, C. (2011). CD155 on Human Vascular Endothelial Cells Attenuates the Acquisition of Effector Functions in CD8 T Cells. Arterioscler. Thromb. Vasc. Biol. 31, 1177–1184.

26. Bensellam, M., Jonas, J., Laybutt, D. (2017). Mechanisms of β-cell dedifferentiation in diabetes: Recent findings and future research directions. J. Endocrinol. 236, JOE–17.

27. Stelzer, G., Rosen, N., Plaschkes, I., Zimmerman, S., Twik, M., Fishilevich, S., Stein, T.I., Nudel, R., Lieder, I., Mazor, Y. et al. (2016). The GeneCards Suite: From Gene Data Mining to Disease Genome Sequence Analyses. Current Protocols in Bioinformatics 54, 1.30.1–1.30.33.

28. Walaszczyk, E., Luijten, M., Spijkerman, A.M.W., Bonder, M.J., Lutgers, H.L., Snieder, H., Wolffenbuttel, B.H.R., van Vliet-Ostaptchouk, J.V. (2018). DNA methylation markers associated with type 2 diabetes, fasting glucose and HbA1c levels: a systematic review and replication in a case–control sample of the Lifelines study. Diabetologia 61, 354–368.

29. Bacos, K., Gillberg, L., Volkov, P., Olsson, A.H., Hansen, T., Pedersen, O., Gjesing, A.P., Eiberg, H., Tuomi, T., Almgren, P. et al. (2016). Blood-based biomarkers of age-associated epigenetic changes in human islets associate with insulin secretion and diabetes. Nature Communications 7, 11089.

30. American, D.A. (2012). Diagnosis and classification of diabetes mellitus. Diabetes Care 35 Suppl 1, S64–S71.

31. Rutter, G. (2010). Think zinc: New roles for zinc in the control of insulin secretion. Islets 2, 49–50.

32. Pourvali, K., Abbasi, M., Mottaghi, A. (2016). Role of Superoxide Dismutase 2 Gene Ala16Val Polymorphism and Total Antioxidant Capacity in Diabetes and its Complications. Avicenna J Med Biotechnol 8, 48–56.

33. Flannick, J., Thorleifsson, G., Beer, N.L., Jacobs, S.B.R., Grarup, N., Burtt, N.P., Mahajan, A., Fuchsberger, C., Atzmon, G., Benediktsson, R. et al. (2014). Loss-of-function mutations in SLC30A8 protect against type 2 diabetes. Nat Genet 46, 357–363.

34. Johansson, B.B., Torsvik, J., Bjørkhaug, L., Vesterhus, M., Ragvin, A., Tjora, E., Fjeld, K., Hoem, D., Johansson, S., Ræder, H. et al. (2011). Diabetes and pancreatic exocrine dysfunction due to mutations in the carboxyl ester lipase gene-maturity onset diabetes of the young (CEL-MODY): a protein misfolding disease. J. Biol. Chem. 286, 34593–34605.

35. Torsvik, J., Johansson, B.B., Dalva, M., Marie, M., Fjeld, K., Johansson, S., Bjørkøy, G., Saraste, J., Njølstad, P.R., Molven, A. (2014). Endocytosis of secreted carboxyl ester lipase in a syndrome of diabetes and pancreatic exocrine dysfunction. J. Biol. Chem. 289, 29097–29111.

36. Newman, A.M., Liu, C.L., Green, M.R., Gentles, A.J., Feng, W., Xu, Y., Hoang, C.D., Diehn, M., Alizadeh, A.A. (2015). Robust enumeration of cell subsets from tissue expression profiles. Nature Methods 12, 453–457.

37. Love, M.I., Huber, W., Anders, S. (2014). Moderated estimation of fold change and dispersion for RNA-seq data with DESeq2. Genome Biol. 15, 550.

